# A CNN model with feature integration for MI EEG subject classification in BMI

**DOI:** 10.1101/2022.01.05.475058

**Authors:** Arunabha M. Roy

## Abstract

**Objective:** Electroencephalogram (EEG) based motor imagery (MI) classification is an important aspect in brain-machine interfaces (BMIs) which bridges between neural system and computer devices decoding brain signals into recognizable machine commands. However, the MI classification task is challenging due to inherent complex properties, inter-subject variability, and low signal-to-noise ratio (SNR) of EEG signals. To overcome the above-mentioned issues, the current work proposes an efficient multi-scale convolutional neural network (MS-CNN).

**Approach:** In the framework, discriminant user-specific features have been extracted and integrated to improve the accuracy and performance of the CNN classifier. Additionally, different data augmentation methods have been implemented to further improve the accuracy and robustness of the model.

**Main results:** The model achieves an average classification accuracy of 93.74% and Cohen’s kappa-coefficient of 0.92 on the BCI competition IV2b dataset outperforming several baseline and current state-of-the-art EEG-based MI classification models.

**Significance:** The proposed algorithm effectively addresses the shortcoming of existing CNN-based EEG-MI classification models and significantly improves the classification accuracy.

## 1. Introduction

Brain–computer interfaces (BCI) or brain-machine interface (BMI) enable a direct communication between the brain and external devices [1, 2, 3] which allows rehabilitation of neuromotor disorders [4], robotic control [5, 6, 7], speech communication [8, 9] etc. In general, BMIs can be invasive, non-invasive or synthetic telepathy [2]. Non-invasive method includes electroencephalography (EEG), functional magnetic resonance imaging (fMRI), magnetoencephalography (MEG), and near-infrared spectroscopy (NIRS) [2, 3]. However, non-invasive BMI through EEG signal collecting through electrodes placed on the scalp has been a popular choice due to its fine temporal resolution, low cost, and user-friendly communication with other electronic devices. Compared with other types of brain signals, EEG has some distinct characteristics such as uniqueness, non-linearity, and non-stationary behaviors which vary with the human brain and the mental state of the particular subjects [10]. Additionally, due to the presence of noise from different muscle artifacts, it poses a challenge to effectively improve the signal-to-noise ratio (SNR) to enhance accuracy in subject classification. Thus, the feature extraction and classification of EEG signals is an important aspect of designing a robust BMI system. The commonly used EEG signals include motor imagery (MI) related mu/beta rhythm (de)synchronization, event-related P300 potentials, and steady-state visually evoked potentials (SSVEPs) [11, 12]. Among these, MI is the most popular in various EEG-based BCI applications [2, 3]. The general workflow of a typical EEG-based MI BCI system is shown in Fig.1 which generally consist of four phases including brain signal acquisition, feature extraction, feature classification, and device control interface. For feature extraction in time-frequency spectrum, wavelet [13] or short-time Fourier-transformation (STFT) [14] have been utilized. Due to the limitation of feature extraction in the same frequency band, the classification accuracy may fall for different subjects. To overcome this, wavelet packet decomposition (WPD) and dynamic frequency feature selection (DFFS) [15] have been employed to obtain better time-frequency features for each subject [16]. However, the procedure is time-consuming and can not be generalized. In regard to feature extraction of EEG signal in space domain, common spatial pattern (CSP) [17], filter bank CSP [18] have shown to be effective in improving accuracy, however, the performance depends on a specific frequency band and does not consider full time-domain feature extraction from different subjects.

**Figure 1:**
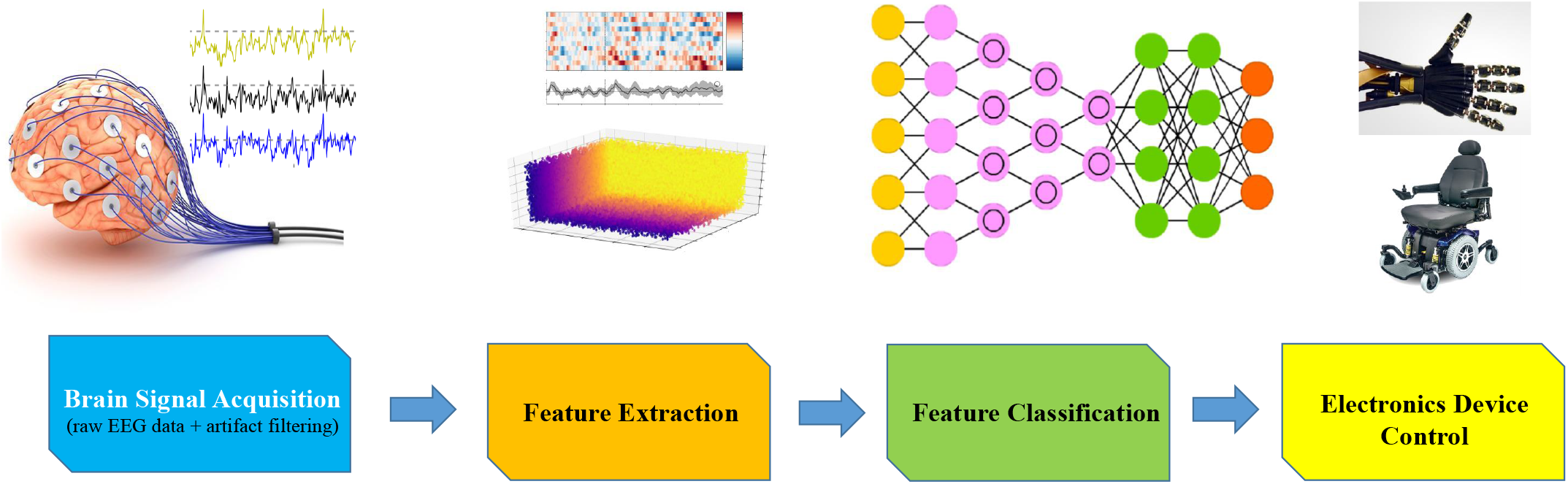
The general workflow of a typical EEG-based MI-BCI system consist of brain signal acquisition, feature extraction, feature classification, and device control interface.

With the advancement of deep learning (DL) in recent years, it illustrates superior performance in MI-BCI classification compared to traditional ML methodologies [14] due to the capability of adapting non-linear and non-stationary signals and extracting important feature information [19] from EEG signal automatically. In this regard, there are several studies have been geared towards the EEG signal classification employing DL, in particular, convolutional neural network (CNN) [14, 20, 21, 22, 23, 24, 25]. A DL model with the combination of CNN and stacked autoencoders (SAE) have been developed which demonstrated the improvement of recognition accuracy for EEG signal classification [14]. In [23], an end-to-end CNN has been designed for efficient MI-EEG signal classification. Some traditional feature extraction methodologies such as wavelet transform (WT) from time-frequency images have been incorporated in CNN for subject classification [26]. A DNN Scheme based on restricted Boltzmann machines (RBM) for MI classification has been proposed in [27]. In addition, EEGNet framework [28] based on compact CNN has been proposed for MI and P300 visual-evoked potentials which demonstrated improvement of classification accuracy compared to the state-of-the-art methods. Along similar line, a deep transfer CNN framework based on the VGG-16 CNN model pre-trained on the ImageNet and a target CNN model for MI EEG signal classification has been proposed in [29]. Furthermore, a 1-D multi-scale CNN [30] based on conditional empirical mode decomposition (CEMD) has been developed which correlates the original EEG signal and intrinsic modal component (IMF) to encode event-related synchronization/de-synchronization (ERS/ERD) information between the channels achieving higher accuracy for EEG signals classification. Additionally, a multiple bandwidth method with optimized CNN framework [31] has been designed for BMI classification with EEG-fNIRS signals. More recently, a hybrid-scale CNN architecture [32] for EEG MI classification has been proposed which demonstrates significant improvement in classification accuracy.

Although, the CNN-based models have achieved better results, there are several issues which cause to hinder the accuracy and performance of the classifier for EEG MI classification. Firstly, most of the CNN-based models consider only a single convolution scale which is not sufficient to extract distinguishable features of several non-overlapping canonical frequency bands of EEG signal efficiently. Secondly, intrinsic feature extraction of the input signal is often ignored which limits CNN’s ability to learn more semantic features from the raw EEG data. Moreover, feature extraction has not been designed to fully integrate into the DL workflow which is the main bottleneck for the deployment of real-time BCI applications with high classification accuracy. Thirdly, one of the common issues of CNN-based models is the lack of sufficient training data which restrain to achieve high classification accuracy for EEG-based MI-BCI classifier.

In order to address the aforementioned challenges and shortcomings, an efficient multi-scale convolutional neural network (MS-CNN) has been proposed for EEG-based MI classification. In the model, a multi-scale convolution block (MSCB) with different convolutional kernel sizes has been designed to extract the effective features of EEG signals from multiple scales for four different frequency bands *δ*, *θ*, *α*, and *β* from original EEG data for MI classification. Moreover, important intrinsic and user-specific features including differential entropy (DE) and neural power spectra (NPS) characteristics have been extracted from the original EEG data and integrated into the proposed algorithm to improve the accuracy and performance of the model. Additionally, different data augmentation (DA) methods such as Gaussian noise (GN), signal segmentation and recombination (S & R), window slicing (WS), and window wrapping (WW) have been employed to further improve the accuracy and robustness of the proposed classifier by increasing training EEG data. It has been found that the current algorithm significantly outperforms several baselines and the current state-of-the-art EEG-based MI classification models with an average classification accuracy of 93.74% on BCI Competition IV2b dataset. Current study provides an effective and efficient framework for designing high performance real-time MI-BCI system.

## 2. Dataset and description of experimental method

In the present work, BCI competition IV 2b dataset [33] has been utilized to evaluate the efficiency and accuracy of the proposed MS-CNN model. The 2b datasets contain EEG data collected from nine healthy subjects. For each subject, the EEG signal data has been recorded and collected from three bipolar EEG channel electrodes with a sampling frequency of 250 Hz. These electrodes (i.e., *C*_3_, *C_Z_*, and *C*_4_) have been positioned according to standardized international 10-20 electrode system [34] as shown in Fig. 2-(a). In order to eliminate the power line signal noise, a band-pass filter allowing EEG signal frequency between 0.5 Hz and 100 Hz with a notch filter at 50Hz was implemented. Additionally, the electrooculogram (EOG) has been recorded using three monopolar electrodes [35, 36]. In the dateset, two types of MI classification tasks has been performed by each subject which include left (class 1) and right-hand movement (class 2) imagination. From the given dataset, there is a total of 5 sessions were considered for each of the subjects. For each subject case, the first two sessions consisting of 120 trials were collected without feedback in EEG signal data. Whereas, the remaining three sessions of 160 trials were recorded with EEG feedback. The schematic of each trial and corresponding timing stamp has been depicted in Fig. 2-(b). For example, consider the first two sessions, the successive events are as following order: at the start of each trial (i.e., t=0) a fixation cross appears; followed by a cue in the form of an arrow at t=3 s to t=4.25 s indicating two distinct MI tasks at once; finally, the subject performs left and right-hand movement imagination according to arrow direction from t=4 s to t=7 s [35, 36, 37].

**Figure 2:**
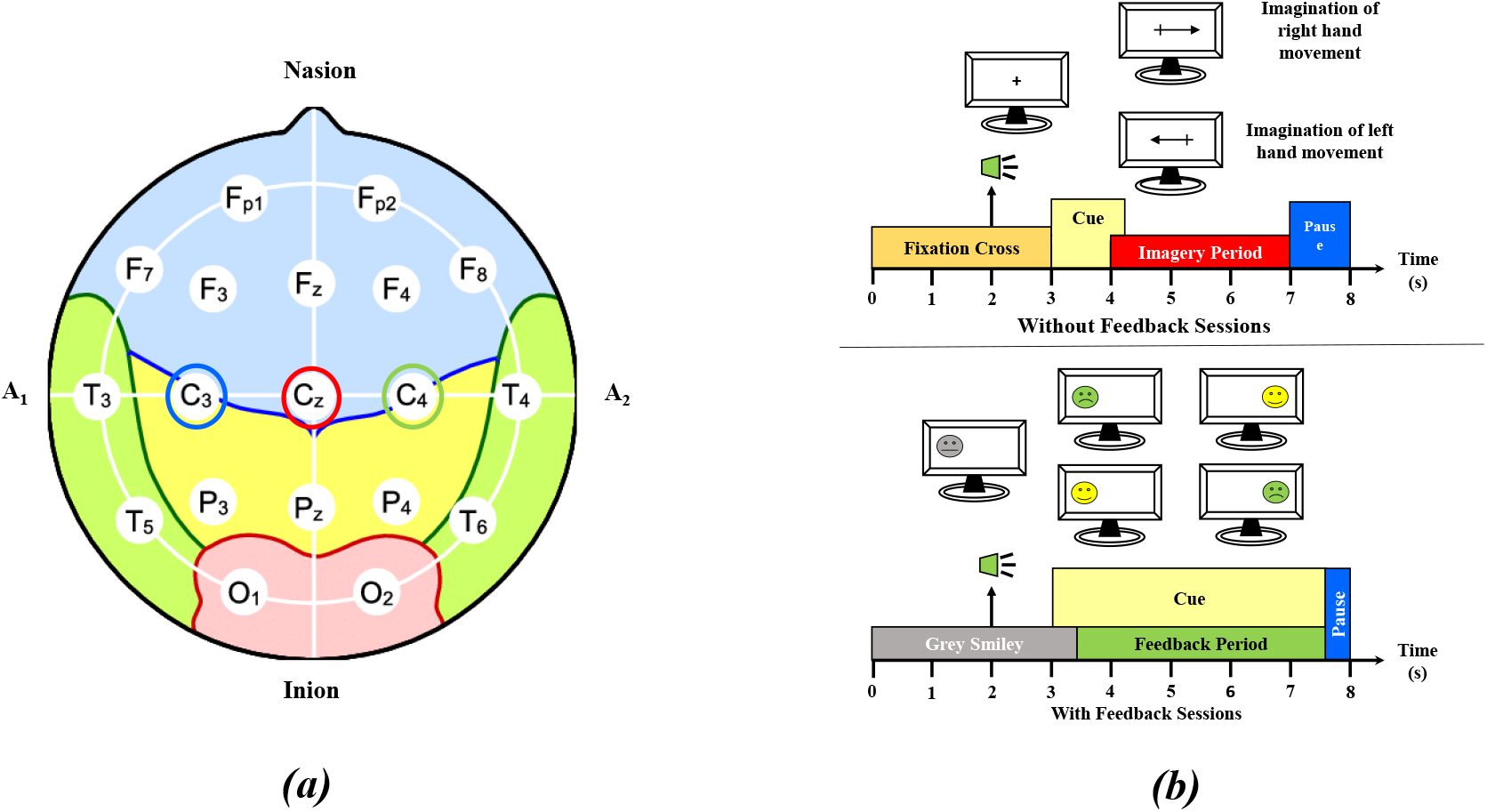
(a) Schematics of electrodes positioning of *C*_3_, *C_Z_*, and *C*_4_ in standardized international 10-20 electrode system; (b) timing scheme of each trial including first two sessions (top) and remaining three sessions (bottom).

**Figure 3:**
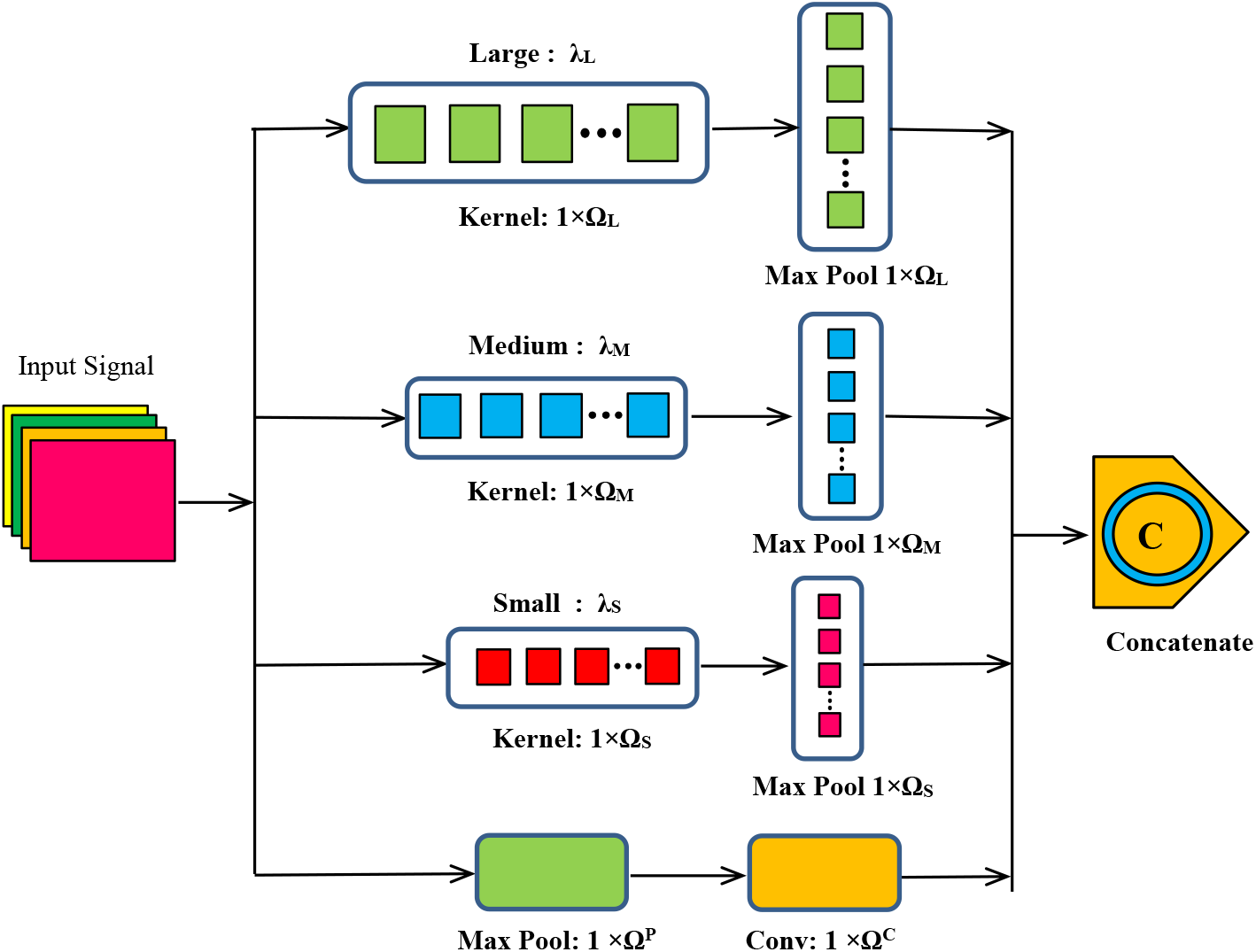
Generalized schematic of network structure for proposed multiscale convolution block (MSCB) consisting of three different convolution scale: large (λ_*L*_), medium (λ_*M*_), and small (λ_*S*_) for multi-scale feature extraction.

## 3. Proposed CNN model

In recent years, CNN has demonstrated significant performance improvement outperforming traditional ML approaches [38] in various applications such as object detection and computer vision [39, 40]. The MI subject classification from the raw EEG signal with high variability and non-stationary noise is a challenging task. In this regard, CNN can be an effective method for extracting the most relevant features and learning the hierarchical representations of high dimensional EEG time series data. CNN is a feed-forward network that is usually comprised of the following components: input layer, convolution (Conv) layer, pooling (Pool) layer, fully connected (FC) layer, and output layer. Generally, the CNN network consists of alternating convolution and pooling layers for extracting features, and a fully connected layer at the end for final classification. Mathematically, the convolution process can be expressed as:

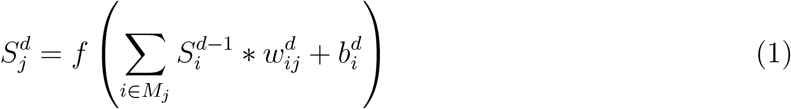

Where 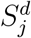 is the *j^th^* feature information in the *d^th^* Conv; 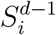 and 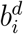 are the *i^th^* feature map and bias term corresponding to *d* – 1^*th*^ and *d^th^* Conv, respectively; 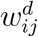 represents connecting weight between *i^th^* feature of the *d* – 1^*th*^ layer and *j^th^* feature of the *d^th^* layer; * denotes convolution operator; *M_j_* represents input feature collection; *f*(●) represents the activation function which can be hyperbolic tangent *f*(*x*) = *tanh*(*x*), sigmoid type *f*(*x*) = 1/(1 + *e*^−*x*^), or rectified linear units (ReLU) *f*(*x*) = *max*(0, *x*) etc. The main functionality of pooling layer is to reduce the spatial size of the representation and the number of network parameters while preserving important and relevant features information. The fully connected layer at the end of the CNN transforms the high dimensional feature map obtained from the previous layer into 1D array. Finally, it is connected to the Softmax layer to predict the classification result. However, in EEG signals, there are several non-overlapping canonical frequency bands corresponds to various distinct behavioral state[41, 42]. Each frequency pattern represents a qualitative assessment of awareness during MI tasks. The low frequency *δ*-bands (1-4 HZ) was found to carry significant class-related information [2, 43, 44]. Additionally, in movement-related MI-BCI systems, *α* (8-13 HZ)and *β* (13-30) rhythms are important due to their high temporal resolution [45]. An increase and decrease of power spectrum in the *β* and *α*-bands results in event-related synchronization (ERS) and event-related desynchronization (ERD), respectively [46, 47]. Recently, it has been revealed that *θ*-band (4-8 Hz) significantly differs between the left/right-hand MI tasks which plays an important role in MI-BCI classification process [32, 48, 49]. Thus, in the current study, we have considered four non-overlapping frequency bands including *δ*, *θ*, *α*, and *β* for feature extraction from original EEG data in our proposed CNN model. A filter bank of 1-4 Hz, 4-8 Hz, 8-13 Hz, and 13-30 Hz has been employed to extract EEG signal information in corresponding frequency bands.

### 3.1 multi-scale convolution block (MSCB)

In the proposed MSCB network, convolution block λ_*L*_ has a relatively large convolution kernel size 1 × Ω_*L*_ which can capture the overall feature map of the EEG signal. Relatively medium convolution kernel size 1 × Ω_*M*_ in convolution block λ_*M*_ can preserve relatively coarse grain feature information. Finally, convolution block λ_*M*_ with small kernel size 1 × Ω_*S*_ can efficiently collect fine-grain localize information.

### 3.2 Multi-scale CNN (MS-CNN)

The proposed multi-scale CNN architecture has been presented in Fig. 4. At first, the inputted EEG signal has been divided into four different frequency bands channels and passed through corresponding MSCB blocks (i.e., MSCB_*i*_, *i* = *δ*, *θ*, *α*, and *β*) to obtain multi-scale features from the EEG signal as shown in Fig. 4. These MSCB blocks have been connected to convolution and then max-pooling layers. The kernel size of 1 × Ω*^CL^* in convolution layer with stride *S_CL_* = 1 can preserve relatively coarse fine-grain feature information. The subsequent max-pooling layer reduces the size of feature information utilizing filter size of Ω^*MP*^ with stride *S_PL_* = 5. The extracted features from the max-pooling layer have been passed through the first flatten layer for feature fusion. During feature fusion, 2D feature vectors are transformed to 1D feature by row concatenation process for all four brunches as shown in Fig. 4. Finally, further concatenation operation transforms all the 1D features from the four branches into a 1D array which has been used as the input to the fully connected layer with number of hidden units *N_h_*. The output of the network is obtained by utilizing the Softmax layer that performs multi-class classification of EEG signals. The proposed network utilizes ReLU as the primary activation function which accelerates the optimization process of the MS-CNN network and increases the classification accuracy for MI-BCI application [30]. Moreover, cross-entropy has been used as a loss function to optimize the model during training. The cross-entropy *L_CE_* for *n* classes can be defined as:

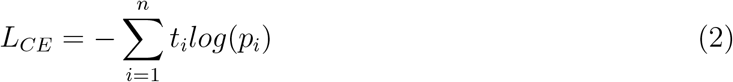

**Figure 4:**
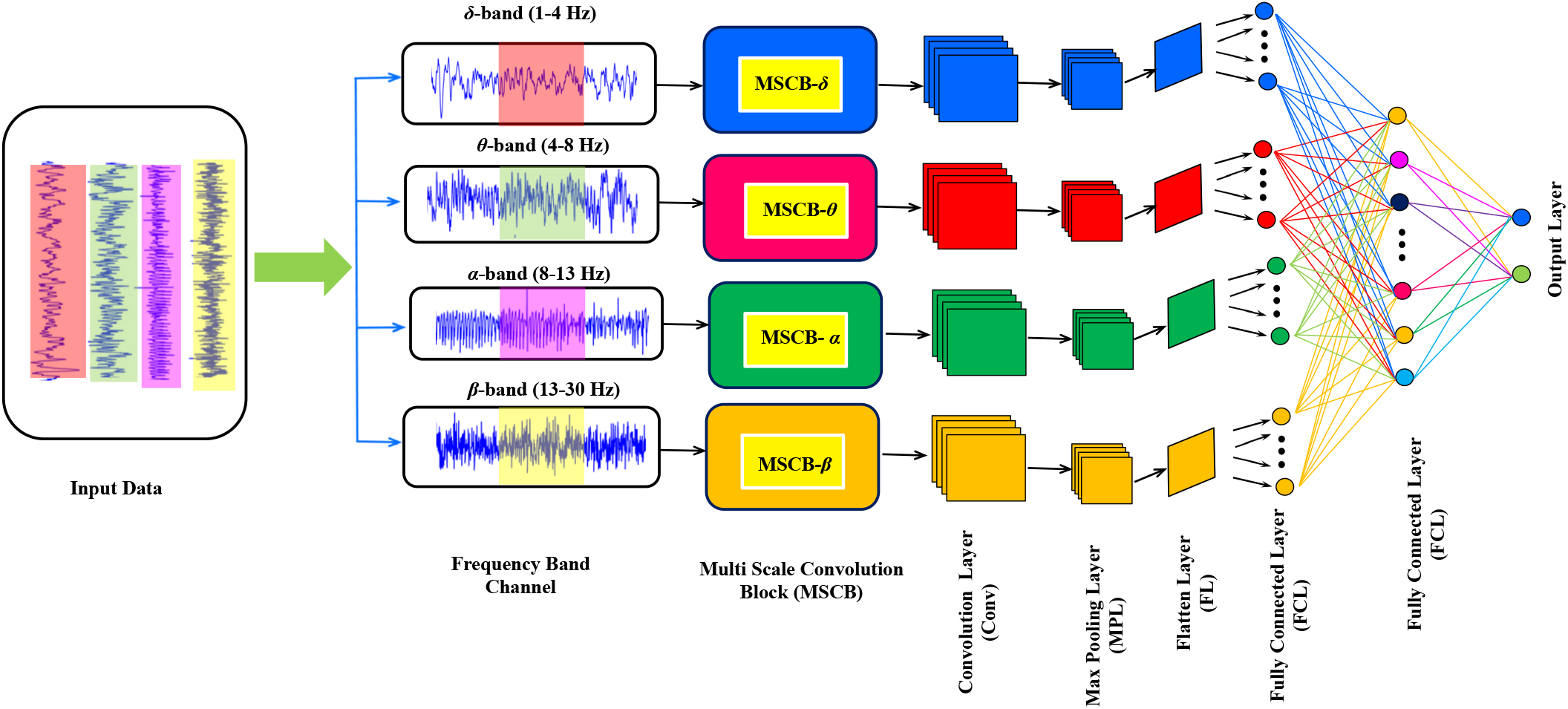
The schematic of proposed multi-scale CNN (MS-CNN) network architecture consisting of four different MSCB as shown in Fig. 3 for efficient multi-scale feature extraction for different non-overlapping frequency bands.

Where *t_i_* is the truth label; *p_i_* is the Softmax probability for *i^th^* class. In the proposed model, the trial matrix size of 1 × *N_L_s__* (where *N_L_s__* is the numeber of data points of each EEG trials) from EEG signal data has been inputted in the MS-CNN; where *L_s_* is the segment of EEG signal in interest and *N_c_* is the number of EEG channels. The time period of a MI trial ranges from 3.5s to 7.5s as shown in Fig. 2-(b). Thus, with the sampling frequency of 250 Hz, we obtain a trial of *L_s_* = 4*s* with *N_L_s__* = 1000 data points.

## 4. EEG Feature Extraction

Due to inter-subject variability of the EEG signals, the accuracy of the classifier often diminish. Thus, it is critical to extract the user-specific features information from EEG data to improve classification accuracy in MI based BCIs. Although there are many features such as statistical, time-domain, frequency-domain, wavelet, auto-regressive coefficients can be extracted from EEG signals [52], however, in the proposed framework, two highly discriminant user-specific features have been extracted and integrated in the MS-CNN model which improve the accuracy and performance of the model.

### 4.1 Differential entropy

From the MI-EEG signal data segment, differential entropy (DE) feature has been extracted which demonstrated superior performance compared to commonly used features [53]. The DE of a given EEG signal *X* satisfying the Gauss distribution *N*(*μ*, *σ*^2^) can be expressed as

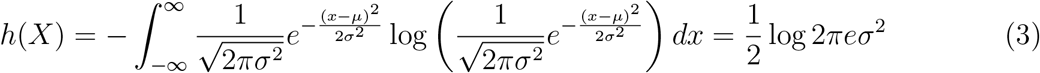

Where *σ* is the standard deviation, *ζ* is the mean value. For a particular EEG signal segment, DE feature is equivalent to the logarithm energy spectrum in a distinct frequency band. Thus, a bandpass filter has been employed to extract four different frequency bands *δ* (1-4 Hz), *θ* (4–8 Hz), *α* (8–13 Hz), and *β* (13-30 Hz) from each EEG channel. Subsequently, short-time Fourier transform has been employed with a non-overlapped hamming window of 4 s on trial EEG signal segment consist of 1000 data points to obtain the energy spectrum of each specified frequency band. Finally, the DE feature has been extracted by calculating the logarithm energy spectrum for each of the aforementioned four frequency bands from single channel. After feature extraction, the shape of the DE feature matrix for each selected EEG segments has been concatenated with MS-CNN main feature matrix.

### 4.2 Neural power spectra

Typically, EEG signal data has been analyzed using only canonically defined frequency bands, ignoring the aperiodic component which may compromise physiological interpretations. However, the EEG neural data contains both periodic and aperiodic components [42]. Recently, the neural power spectra (NPS) model has been introduced which combines both aperiodic component and putative periodic oscillatory peaks [42]. In this model, the neural PSD has been characterized by the power, specific center frequency, and bandwidth without requiring predefining specific EEG frequency bands and controlling for the aperiodic component. Additionally, the characteristics of these aperiodic components allow one to measure and compare the 1/*f*-like components between inter-subject variability of the EEG signals. This model has been utilized to extract periodic and aperiodic features of EEG data in the present study. To measure the periodic activity, the power relative to the aperiodic component has been calculated with each peak can be described by Gaussian in terms of parameters *a*, *c* and *w*, where *a* is the height of the peak, over and above the aperiodic component; *c* is the center frequency of the peak; *w* is the width of the peak; *F* is the array of frequency values. Each Gaussian, *n* referred to as *G*(*F*)_*n*_ can be expressed as [42]:

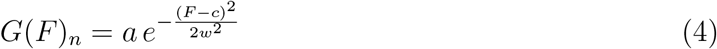

Whereas, the aperiodic activity component without any characteristic frequency can be expressed as the function *L*(*F*) as follows:

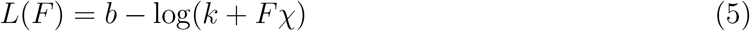

Where *b* is the broadband offset; *χ* is the exponent of the aperiodic fit; *k* is the ‘knee’. Finally, across a set of frequencies *F*, NPS can be expressed as *NPS*(*F*) = *G*(*F*)_*n*_ + *L*(*F*). For better clarity, an example of NPS containing a strong peak in *α* band with overlapping nature of periodic and aperiodic spectral features for two classes have been shown in Fig. 5-(a, b). Consequently, the power ratio (spectral power in the bin normalized by power in all spectral bins) for four different canonical frequency range from the electrodes *C*_3_, *C_Z_*, and *C*_4_ reveals that the α range power decreases as shown in Fig. 5-(c, d, e) on the opposite (i.e. contralateral) hemisphere when MI on one side is performed for a particular class. The apparent differences between the different electrode’s PSDs, in particular, relative PSDs of *C*_3_ (see Fig. 5-c) and *C*_4_ (see Fig. 5-e) channels vary between the two MI classes. Clearly, NPS is a highly discriminant user-specific feature that improves the accuracy and performance of the model significantly (see section 6.3). In the present study, NPS features containing periodic (*c*, *w*, and *P_band_*) and aperiodic parameters (*b*, *k*, and *χ*) have been extracted for each of the four frequency bands for each channel and finally concatenated with MS-CNN main feature matrix as shown in Fig. 8.

**Figure 5:**
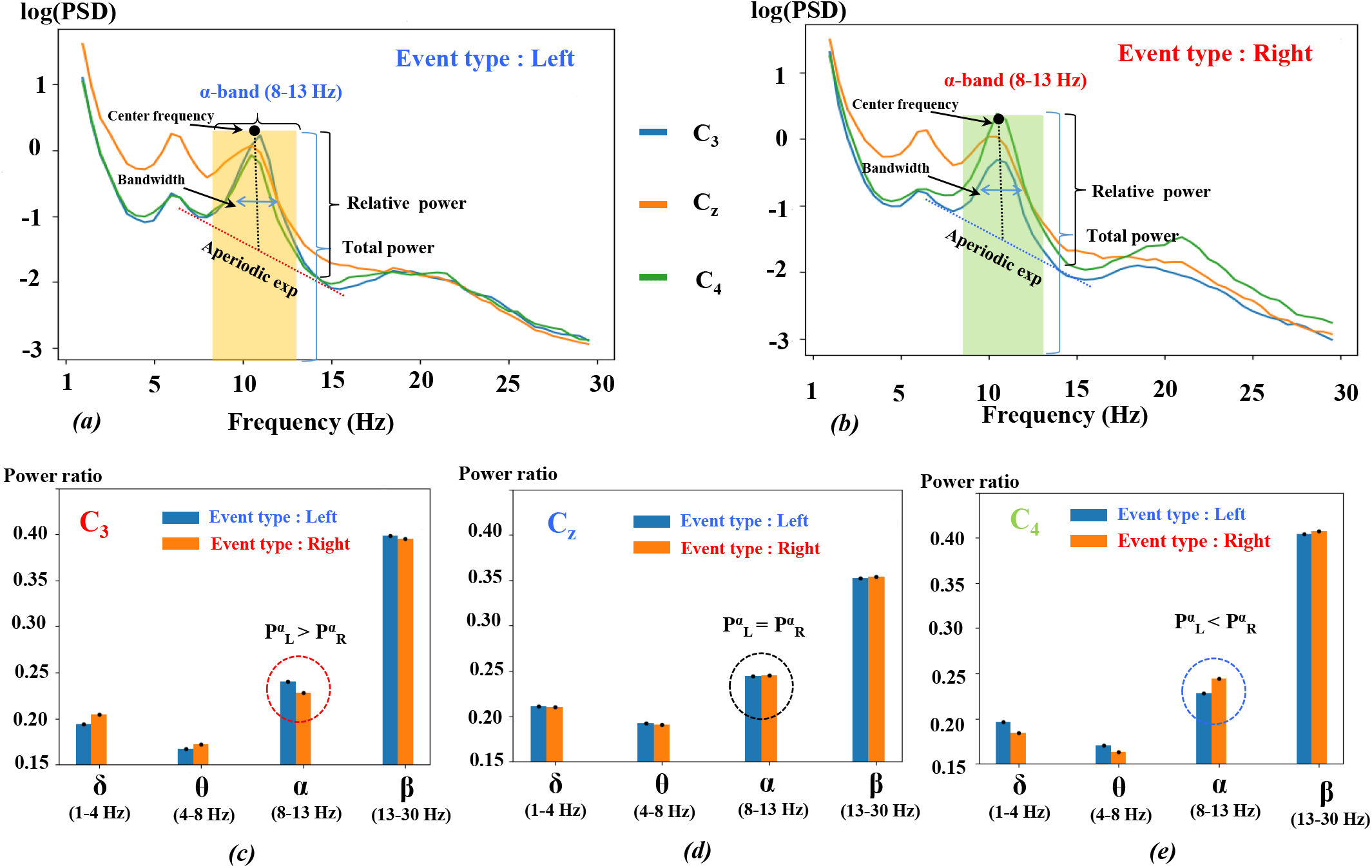
(a, b) NPS containing strong peak in *α* band (8-13 Hz, yellow and green region) and secondary *θ* (not marked) with overlapping nature of periodic and aperiodic spectral features for left and right event type, respectively; (c, d, e) comparison of power ratio in left and right event type for four different canonical frequency range from the electrodes *C*_3_, *C_Z_*, and *C*_4_.

## 5. Data augmentation of EEG signal

In DL based MI-BCI system, the accuracy of the MI classifier is greatly dependent on the volume of EEG training data. Without sufficient data, the accuracy may drop. Hence, to improve the accuracy and robustness of the CNN classifier, data augmentation (DA) methods can be employed to generate new sample data from existing EEG training sample [54]. However, improper DA might lead to a decrease in the performance of the classifier. In the present study, four different types of DA methods have been chosen and implemented specifically for EEG-based MI-BCI systems to increase the volume of EEG data during training and improve the performance of the classier (see section 6.4).

### 5.1 Gaussian noise

In the first DA method, noise has been added to the original training data. The EEG signal has strong randomness and highly non-stationary characteristics. Thus, randomly added local noise may alter the important EEG feature. In order to preserve the important local feature of EEG data, Gaussian noise (GN) has been added to the original training data [53]. The probability density function *P_G_* of a Gaussian random variable *ξ* can be expressed as follows:

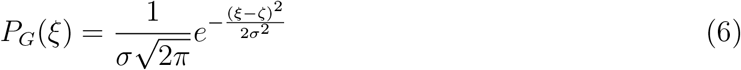

Where *σ* is the standard deviation, *ζ* is the mean value. In the present work, *ζ* = 0 has been prescribed to ensure that the amplitude of the original EEG signal remain unchanged after the addition of noise. During DA, different noise intensity i.e., *σ*=0.001, 0.005, 0.01, 0.05, 0.10, 0.25, and 0.5 have been considered.

### 5.2 Signal segmentation and recombination in time domain

Additionally, EEG data transformation including signal segmentation and recombination (S & R) has been utilized as a second DA method to further expand the dataset. In this procedure, each training EEG trial for the particular class has been subdivided into multiple segments and then new trials are generated by combining segments collected from different and randomly selected trials from the same training class data [55]. If Δ = {*S^i^*}, *i* ∈ [1, *n*] is the set of total *n* number of EEG training data for given class, each training EEG trial *S^i^* for the particular class has been subdivided into *K* consecutive and non-overlapping segments 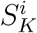 and then generating a new trials 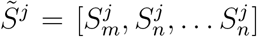 by combining segments from different and randomly selected training trials from the same class. The schematic of S & R - DA procedure has been shown in Fig. 6. Considering the original trials *S_m_* and *S_n_* from the same class have been segmented into 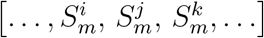 and 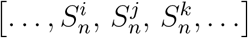. These segments have been recombined in time domain to obtain *N* additional DA trials. In the present work, input EEG trials (i.e. left/right-hand MI) have been considered as same label with each trial has been segmented into 4 division with 250 data points (i.e., 1 sec long segment).

**Figure 6:**
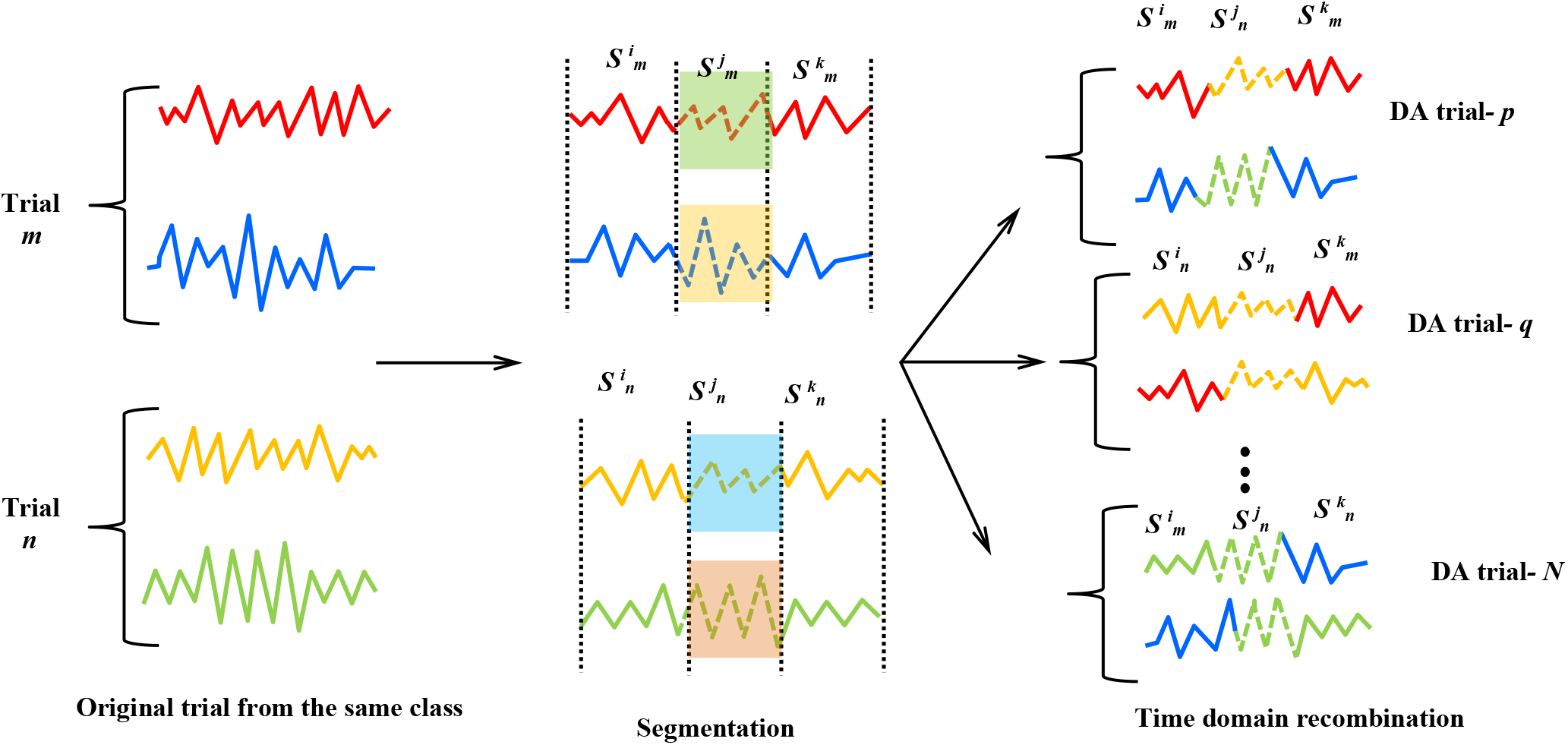
Procedure of EEG signal segmentation and recombination in time domain for data augmentation.

### 5.3 Window slicing

In the window slicing (WS) DA method, EEG time series data have been extracted in slices and classification has been performed at different slice levels [56]. During training, each slice of the corresponding class has been assigned to the same class where the size of the slice is one of the parameters for the DA. In the present work, a window of 90% of the training EEG data has been chosen randomly and interpolated back to the original size to fit with the classifier [56, 57].

### 5.4 Window Warping

In the last DA technique, window warping (WW) [56] DA has been utilized which expands or contracts random windows of the EEG training data by some specific value. Considering the length of the original EEG signal as a parameter, WW warps a randomly selected segment as shown in Fig. 7. Although, WW generates input time series of different lengths, however, the issues can be overcome by performing DA on transformed EEG data having equal lengths [56]. The present study considers a random window of 10% of the original EEG data and wrapped it by speeding it up by 2 or slowing it down by 0.5.

**Figure 7:**
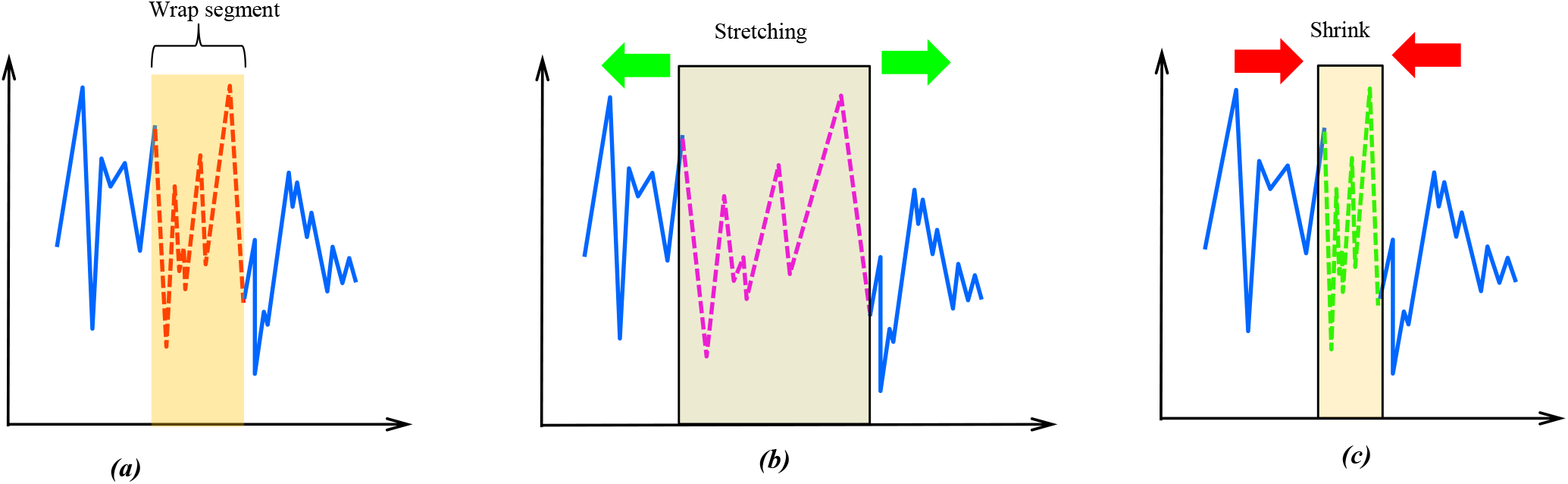
Schematic of window warping data augmentation technique for EEG signal.

## 6. Results and discussion

In this section, the performance and accuracy of the proposed model have been discussed and compared with several existing methods. The BCI competition IV-2b EEG dataset includes 9-subject and 2-class motor-imagery (right hand, left hand) with five sessions for each subject. The first two sessions (identifiers: 01T and 02T) have been used to train the classifier for all models [35]. The third session (identifiers: 03T) has been employed during validation. The last two sessions (identifiers: 04E and 05E) have been strictly utilized for evaluating the corresponding trained classifier [35, 36]. The EEG trial length of *L_s_* = 4*s* with *N_L_s__* = 1000 data points has been used. The proposed MS-CNN model has been fine-tuned on the validation set and then applied to testing sets. The final fine-tuned network parameters 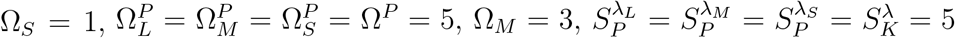 and Ω_*L*_ = 5 have been prescribed. During training, a batch size of 16 has been chosen to optimize the convergence speed and achieve higher classification accuracy. In the proposed model, stochastic gradient descent (SGD) has been utilized as a training strategy with 500 epochs prescribing an exponential decay rate of 0.8 for optimizing the cross-entropy loss function. Due to the high conditionality of the optimization procedure in MI-BCI classification, SGD reduces the computational burden by achieving faster convergence. The initial learning rate is set to 0.001 and the learning rate decays every 30 epochs with an exponential decay rate of 0.5. Lastly, L2 regularization with a regularization parameter value of 0.02 and dropout technique with a dropout probability value of 0.5 (in the first FC layer) have been prescribed to prevent over-fitting. Finally, the models were implemented using the Keras API with TensorFlow as the lower level backend library. Additionally, MNE v0.23 [58], PyEEG [59], NeuroDSP [60], and FOOOF [61] libraries have been utilized for data pre-processing, feature extraction, spectral-domain and NPS analysis. The flowchart representing the overall workflow of the proposed framework has been shown in Fig. 8. For each subject, the experiment has been executed 10 times. The accuracy performance of the proposed framework is evaluated on the prepared testing set. The average classification accuracy and standard error 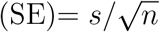 (*s* is the sample standard deviation; *n* is the sample size) have been calculated from the test datasets.

**Figure 8:**
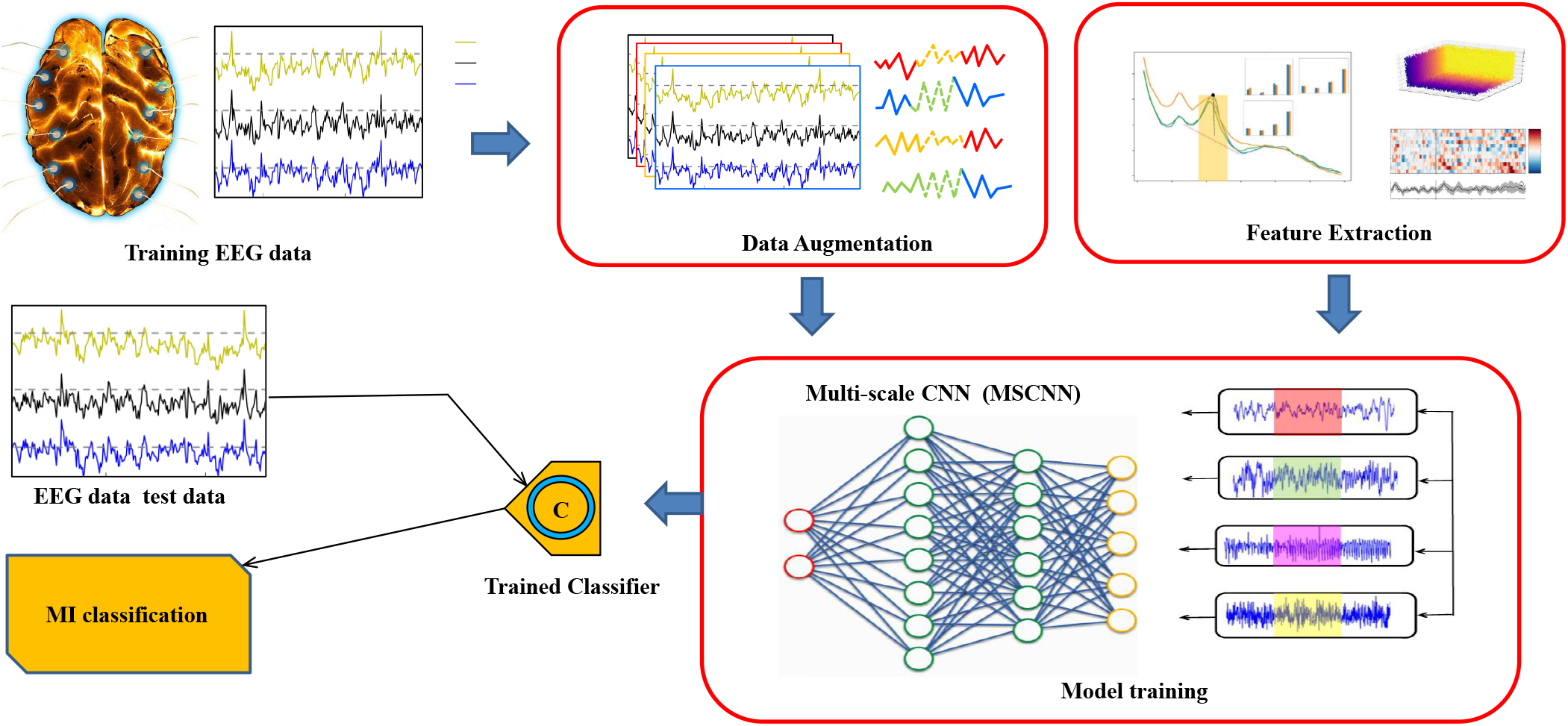
Flowchart representing the overall workflow of the proposed framework for EEG-based MI BCI system based on DA and feature integration.

### 6.1 Performance comparison with different algorithms

In this paper, the average classification accuracy has been used to evaluate different classification models. The global averaged accuracy is defined as the ratio of the number of correctly classified samples to the total number of samples. The performances of the proposed MS-CNN model has been compared with three popular traditional ML models in MI-BCI classification: *K*-nearest neighbor(*K*-NN), linear discriminant analysis (LDA), and support-vector machine (SVM) as the baseline models. Additionally, the standard CNN model has also been considered as the baseline DL model. Note, the standard CNN structure consisting of 4 convolution and max-pooling layers is deeper than the MS-CNN model. For fair comparison, all feature extraction and data augmentation methods have been applied in all aforementioned baseline models. Additionally, model hyper-parameters have prescribed the same as the proposed model. As shown in Table 3, the proposed MS-CNN has obtained the best accuracy across all subjects among all baseline models with an average accuracy of 93.74% for the BCI competition IV-2b dataset. Compared to K-NN, LDA, and SVM, it has achieved d 26.76%, 24.75%, and 26.96% improvement in average classification accuracy, respectively. The proposed model has achieved 16.92% accuracy improvement over the standard CNN model as shown in Fig. 9 indicating the superior performance of MS-CNN. Additionally, the MS-CNN model has the lowest SE value which indicates that the proposed model has the capability of the subject-independent representation of EEG data and better generalization of the classifier compared to other baseline methods. Furthermore, the performance of MS-CNN model on the recognition of two different MI classes (left and right hand) have been measured by precision (*P*), recall (*R*), and F1-score and compared with SVM and standard CNN model as shown in Fig. 10-(a). For binary classification, sample data can be classified into four different categories: true positive (TP), false positive (FP), true negative (TN), and false negative (FN), based on the true class and the model-predicted class. The evaluation matrices *P*, *R*, F1 can be defined as

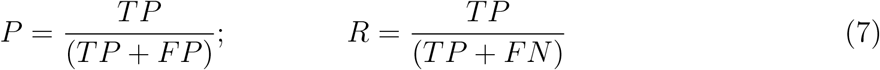

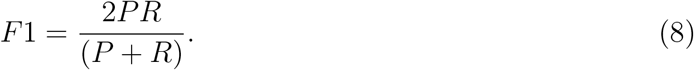

**Table 1:** Main network parameters of generalized multi-scale convolution block (MSCB) as shown in Fig. 3.

| Layer type | Kernel size | No of Kernel | Stride | Padding | Feature map |
| --- | --- | --- | --- | --- | --- |
| Convolution $\lambda_L$ | $\Omega_L^C = 5$ | $N_K^{\lambda_L} = 14$ | $S_K^{\lambda_L} = 1$ | SAME | $L_s \times N_K^{\lambda_L}$ |
| Max Pool $\lambda_L$ | $\Omega_L^P$ | — | $S_P^{\lambda_L}$ | SAME | $(L_s/S_P^{\lambda_L}) \times N_K^{\lambda_L}$ |
| Convolution $\lambda_M$ | $\Omega_M^C = 3$ | $N_K^{\lambda_M} = 14$ | $S_K^{\lambda_M} = 1$ | SAME | $L_s \times N_K^{\lambda_M}$ |
| Max Pool $\lambda_M$ | $\Omega_M^P$ | — | $S_P^{\lambda_M}$ | SAME | $(L_s/S_P^{\lambda_M}) \times N_K^{\lambda_M}$ |
| Convolution $\lambda_S$ | $\Omega_S^C = 1$ | $N_K^{\lambda_S} = 14$ | $S_K^{\lambda_S} = 1$ | SAME | $L_s \times N_K^{\lambda_S}$ |
| Max Pool $\lambda_S$ | $\Omega_S^P$ | — | $S_P^{\lambda_S}$ | SAME | $(L_s/S_P^{\lambda_S}) \times N_K^{\lambda_S}$ |
| Max Pool $\lambda$ | $\Omega^P$ | — | $S_P^\lambda$ | SAME | $(L_s/S_P^\lambda) \times N_f$ |
| Convolution $\lambda$ | $\Omega^C$ | $N_K^\lambda = 24$ | $S_K^\lambda = 1$ | SAME | $(L_s/S_P^\lambda) \times N_K^\lambda$ |
| Concatenate | — | — | — | — | $(L_s/S_P) \times 56$ |

**Table 2:** Main network parameters of proposed multiscale CNN (MS-CNN) model as shown in Fig. 4.

| Layer type | Kernel size | No of Kernel | Stride | Padding | Feature map |
| --- | --- | --- | --- | --- | --- |
| MSCB | - | - | - | - | $(L_s/S_P) \times 56$ |
| CL | $\Omega^{CL}$ | 112 | $S_{CL} = 1$ | SAME | $(L_s/S_P) \times 112$ |
| MP | $\Omega^{MP}$ | - | $S_{PL}$ | SAME | $L_s/(S_P \times S_{PL}) \times 112$ |
| 1<sup>st</sup> FC | - | - | - | - | 400 |
| Dropout layer | - | - | - | - | - |
| 2<sup>nd</sup> FC | - | - | - | - | 300 |
| Output layer | - | - | - | - | 2 |

**Table 3:** Average accuracy values of each subject from *K*-NN, LDA, SVM, CNN, and proposed MS-CNN models where bold indicates the best result from the corresponding model

| Subject | Average Accuracy |  |  |  |  |
| --- | --- | --- | --- | --- | --- |
| | $K$-NN | LDA | SVM | CNN | MS-CNN |
| B01 | 68.32 | 68.77 | 69.45 | 74.52 | **93.45** |
| B02 | 54.39 | 56.46 | 58.92 | 61.29 | **88.71** |
| B03 | 60.77 | 61.29 | 63.89 | 73.67 | **89.14** |
| B04 | 80.22 | 84.49 | 90.22 | 91.45 | **95.37** |
| B05 | 72.17 | 74.32 | 76.23 | 81.43 | **95.11** |
| B06 | 63.89 | 70.35 | 71.47 | 76.89 | **93.82** |
| B07 | 66.32 | 65.60 | 69.68 | 75.89 | **94.70** |
| B08 | 71.17 | 72.12 | 73.41 | 78.83 | **96.81** |
| B09 | 64.97 | 67.55 | 69.77 | 76.82 | **96.73** |
| Avg. Accuracy with SE | 66.98 ±1.67 | 68.99 ±1.49 | 71.45 ±2.01 | 76.76 ±1.84 | **93.74±1.09** |

**Figure 9:**
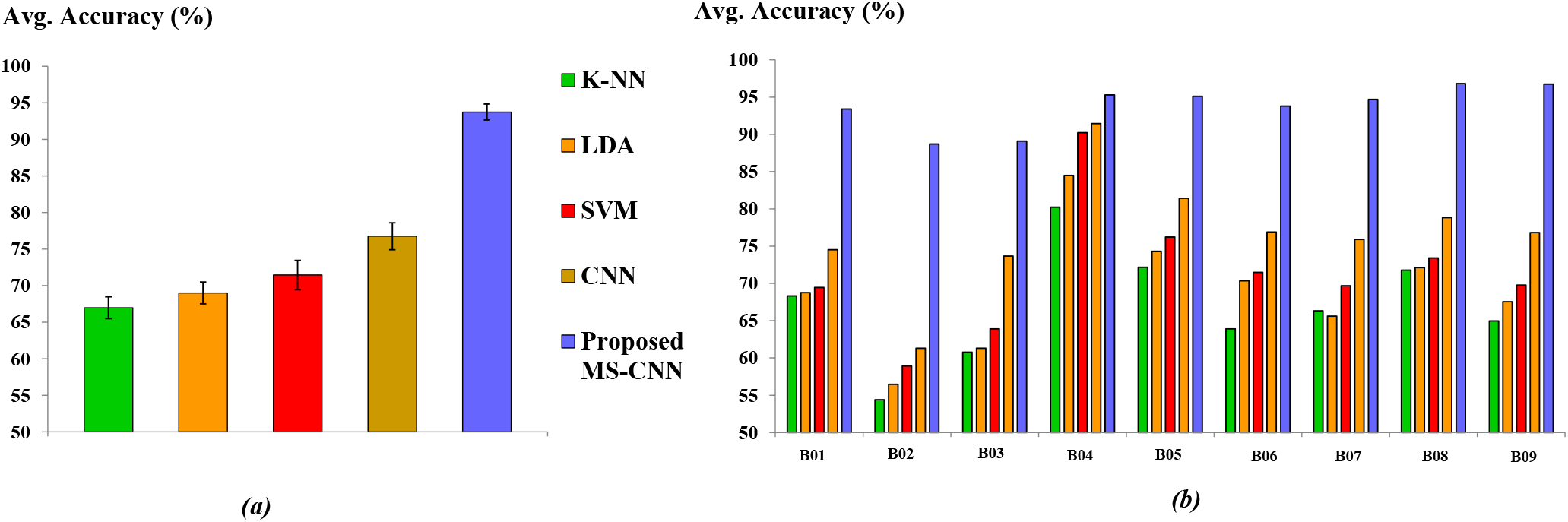
Comparison bar chart of (a) average accuracy with SE (in %) (b) average accuracy of each subject between *K*-NN, LDA, SVM, CNN, and proposed MS-CNN models.

**Figure 10:**
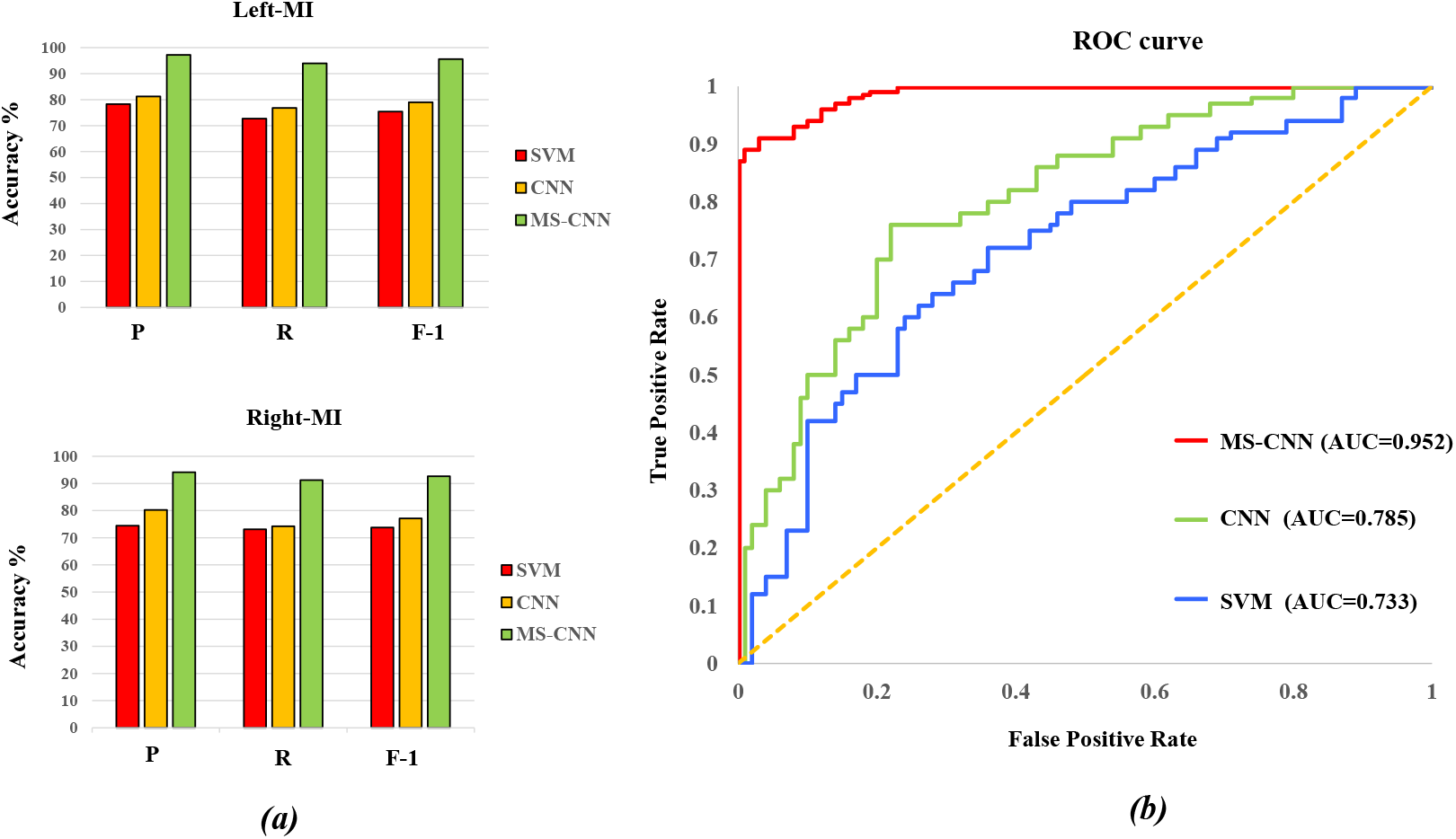
(a) Comparison bar chart of *P*, *R*, F-1 (in %) for the two different MI (left and right hand); (b) ROC curves and corresponding AUC values obtained from SVM, CNN, and MS-CNN models.

The larger values of *P*, *R*, F1 indicate better performance of the model. The proposed MS-CNN model has achieved best *P*, *R*, F1 values of 97.25%, 93.99%, 95.59% and 94.15%, 91.22%, 92.66% for left and right MI, respectively. Finally, ROC (receiver operating characteristic) curve has been plotted from the true positive rate (TPR) or *R* (in the ordinate) and false positive rate (FPR) data (in the abscissa) from SVM, standard CNN, and MS-CNN result as shown in Fig. 10-(b). The area under the ROC curve is expressed in AUC and ranges from 0.5 to 1. The closer the AUC is to 1.0, the higher the performance of the model. As shown in Fig. 10-(b), SVM, standard CNN, and MS-CNN have achieved AUC values of 0.733, 0.785, and 0.952, respectively indicating the best performance of MS-CNN compared to other classification models. The total number of trainable network parameters of MS-CNN is 297,056 compared to 334,236 of the standard CNN. Both models take a similar average time-frame (within 2 *min*) to train, whereas standard CNN takes a slightly higher training time due to a higher number of trainable parameters. The comparison demonstrates that the MS-CNN can have significantly better accuracy and learning capability for different subject classes with less training time. It is noteworthy to mention, both DL-based standard CNN and MS-CNN models demonstrate superior performance in MI classification accuracy than traditional baseline ML models for all subjects. The comparison illustrates the advantages of the DL model for its strong feature extraction capability and capturing abstract-advanced information among different subjects compared to traditional ML methods in EEG-based MI-BCI systems.

### 6.2 Influence of convolution kernel size

In this section, different combinations of the convolution kernel size (i.e, Ω_*S*_, Ω_*M*_, and Ω_*L*_) in MSCB have been studied to explore the influence of kernel sizes on the performance of the model. From the comparison as shown in Fig. 11, it can be shown that with increasing kernel size, in particular, for combinations (Ω_*S*_ = 3, Ω_*M*_ = 7, Ω_*L*_ = 9) and (Ω_*S*_ = 3, Ω_*M*_ = 9, Ω_*L*_ = 11) classification accuracy drops to 89.78% and 87.39%, respectively. However, the combination of relatively small kernel size, classification accuracy, as well as speed, improves. Comparing all the results, it can be found that the best classification result corresponds to (Ω_*S*_ = 1, Ω_*M*_ = 5, Ω_*L*_ = 7) with an accuracy of 93.99%. However, in this study, combinations of Ω_*S*_ =1, Ω_*M*_ = 3, and Ω_*L*_ = 5 has been selected which is computationally more efficient for real-time MI BCI system compromising only 0.25% accuracy compared to result corresponds to (Ω_*S*_ = 1, Ω_*M*_ = 5, Ω_*L*_ = 7). Additionally, the selected combination of kernel scales has achieved the smallest SE value of 1.09%, which is 0.20% lower than the best result. Hence, the chosen combination of kernel sizes in MSCB optimizes the performance of the MS-CNN model in-terms of both classification accuracy and speed.

**Figure 11:**
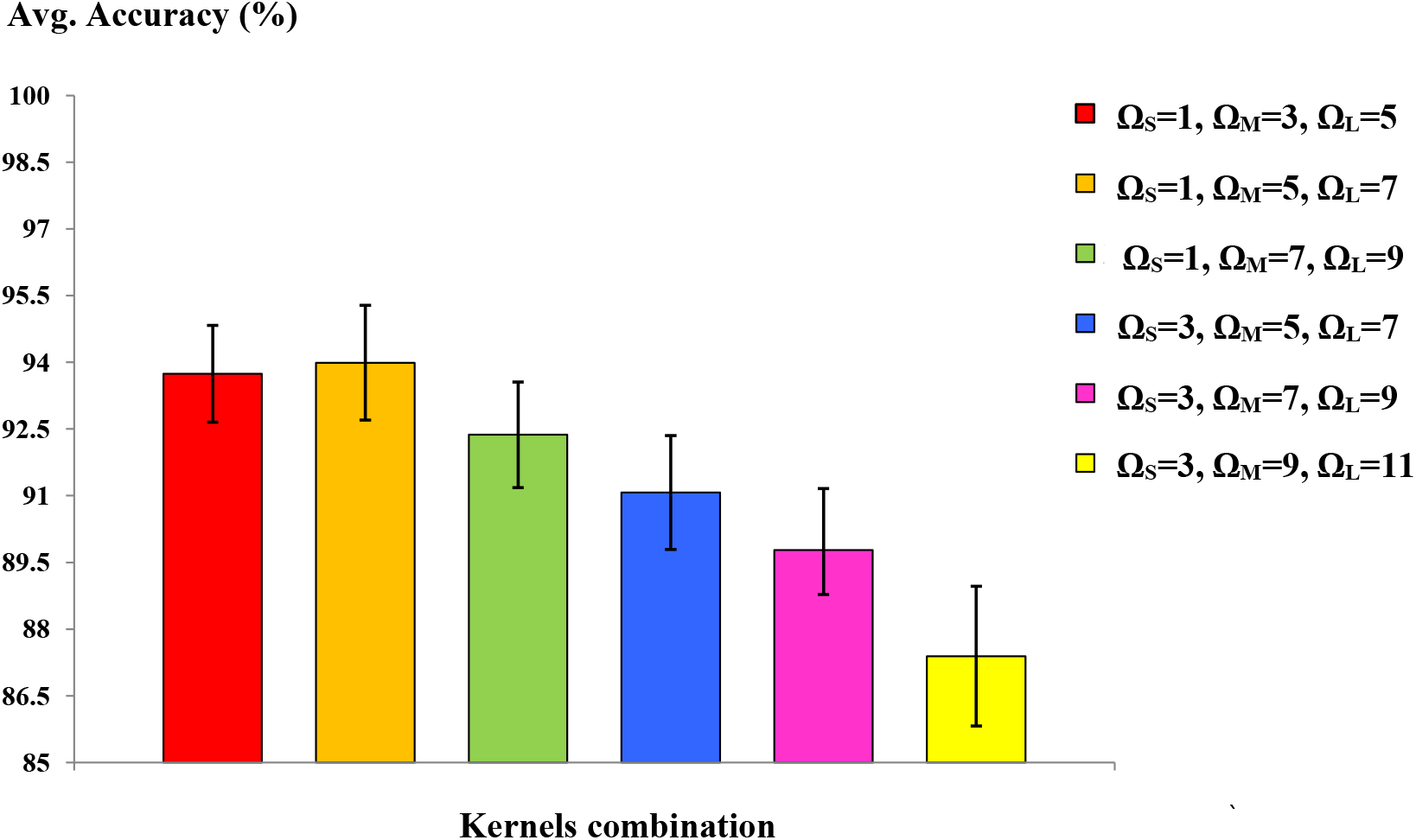
Comparison bar chart of average accuracy with SE (in %) for different combinations of convolutional kernel sizes in multi-scale convolution block (MSCB).

### 6.3 Influence of feature extraction

In this study, the influence of different feature extraction (FE) techniques on classification accuracy has been explored as shown in Table 4. At first, different feature extraction methods that include DE and NPS have been considered individually and the corresponding accuracy of the classifier has been determined. Finally, both the future extraction methods have been employed to obtain the final accuracy of the model. Note, DA methods have not been considered during this study. From the comparison in Fig. 12-(a), it is clear that feature integration significantly improves the accuracy of the model when accuracy increases 10.98% compared to the situation when the proposed model does not utilize any FE techniques. It is noteworthy to mention, the influence of NPS on the performance of the model is significant with 9.21% accuracy gain. From the analysis, it can be inferred that both DE and NPS FEs play an important role for improving overall performance of the MS-CNN model.

**Table 4:** Influence of different intrinsic feature extraction techniques on the average classification accuracy of the proposed MS-CNN model.

| Average Accuracy with SE (%) |  |  |  |
| --- | --- | --- | --- |
| No FE | DE | NPS | All |
| $77.67 \pm 1.89$ | $83.19 \pm 1.67$ | $86.88 \pm 1.42$ | $88.65 \pm 1.24$ |

**Figure 12:**
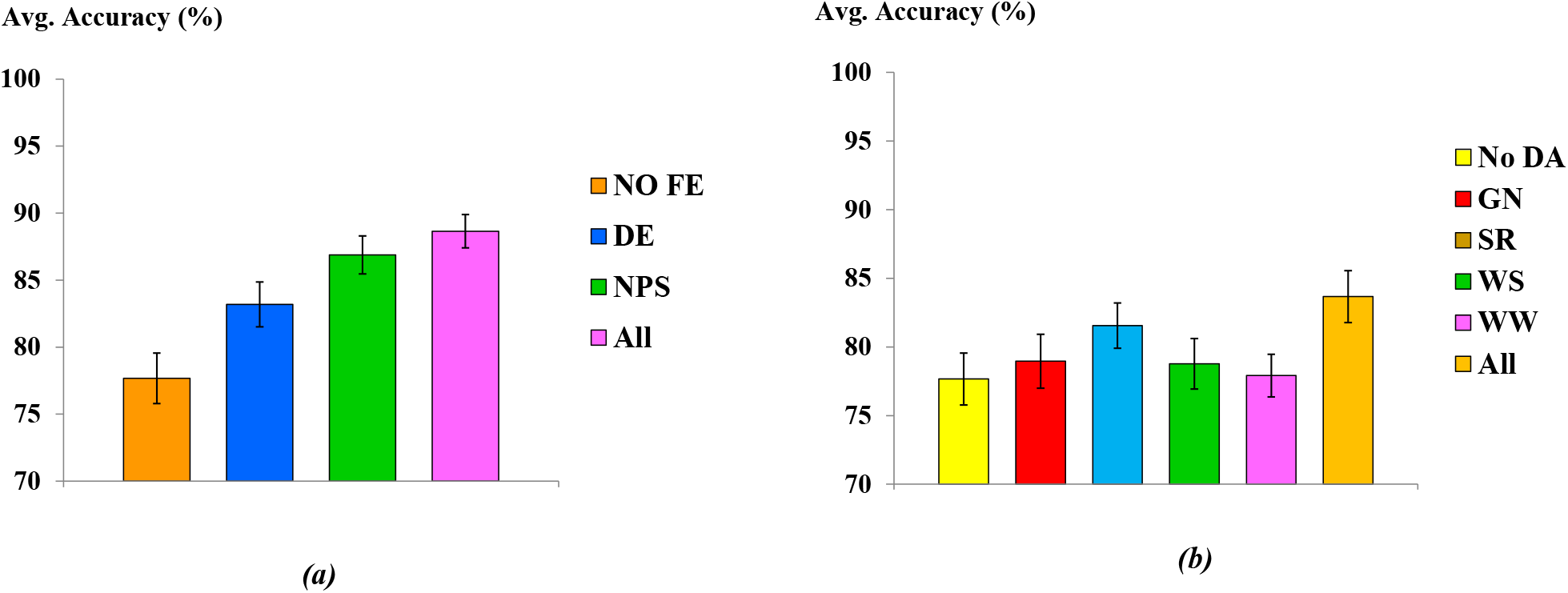
Comparison bar chart of average accuracy with SE (in %) for the influence of different (a) feature extraction techniques; (b) data augmentation methods in the proposed MI-BCI classification framework.

### 6.4 Influence of data augumentation methods

In the section, the influence of different data augmentation (DA) methods including Gaussian noise (GN), segmentation and recombination (S & R), window sliding (WS), and window wrapping (WW) have been evaluated on the performance of the classifier. Note, FE techniques have not been employed during the comparison. As shown in Table 5, the proposed model has reached the accuracy value of 77. 67% without any DA. However, different DA methods improve the accuracy individually, in particular, there is a 3.89% accuracy gain by employing S & R. The combined effect of all DE methods demonstrated an overall 6. 01% accuracy improvement indicating the importance of proposed DA methods for achieving better performance and robustness of the current framework.

**Table 5:** Influence of data augmentation methods on the average classification accuracy of the proposed MS-CNN model.

| Average Accuracy with SE (%) |  |  |  |  |  |
| --- | --- | --- | --- | --- | --- |
| No DA | GN | S & R | WS | WW | All |
| 77.67 ±1.89 | 78.96 ±1.96 | 81.56 ±1.65 | 78.81±1.84 | 77.92±1.55 | 83.68±1.34 |

### 6.5 Comparison with different state-of-the-art ML models

In order to evaluate and compare with the performance of different ML classification models, average Cohen’s kappa-coefficient (*κ*) has been utilized to measure the accuracy of the corresponding classifier. In Table 6, *κ* values obtained from the MS-CNN model have been compared with some of state-of-the-art machine learning models for BCI Competition IV 2b datasets. These ML models have utilized different feature extraction methodologies including CSP [63], filter bank CSP (FBCSP) [64], Hilbert transform (HT) [65], wavelet packet decomposition (WPD) [16], and empirical mode decomposition (EMD) algorithm considering different classification methods such as SVM [64, 66], least squares twin SVM (LSTSVM) [63], LDA [65], and K-NN [16]. From the overall comparison, it can be seen that the proposed MS-CNN model outshines other ML models significantly in terms of *κ* values for all nine subjects, as shown in Fig. 13. Comparing average *κ*, MS-CNN has achieved the highest *κ* value of 0.92 which is 22.2% and 26.01% improvement over the state-of-the-art ML methods in [16] and [64], respectively.

**Table 6:** Comparison of the *κ* values for different subjects with existing state-of-the-art ML models with bold indicates the best result from the corresponding model.

| Method | Subject |  |  |  |  |  |  |  |  |  |
| --- | --- | --- | --- | --- | --- | --- | --- | --- | --- | --- |
|  | B01 | B02 | B03 | B04 | B05 | B06 | B07 | B08 | B09 | Avg |
| CSP+EMD-SVM[66] | 0.44 | 0.35 | 0.32 | 0.60 | 0.33 | 0.32 | 0.51 | 0.61 | 0.37 | 0.42 |
| CSP-PSO based LSTSVM [63] | 0.31 | 0.17 | 0.19 | 0.88 | 0.47 | 0.58 | 0.63 | 0.71 | 0.54 | 0.50 |
| HT-SVM, LDA [65] | 0.61 | 0.31 | 0.32 | 0.99 | 0.78 | 0.72 | 0.63 | 0.78 | 0.74 | 0.65 |
| FBCSP-SVM[64] | 0.48 | 0.29 | 0.25 | 0.97 | 0.98 | 0.70 | 0.67 | 0.84 | 0.79 | 0.66 |
| WPD+SE-Isomap K-NN[16] | 0.69 | 0.33 | 0.26 | 0.92 | 0.78 | 0.96 | 0.64 | 0.73 | 0.94 | 0.70 |
| **Proposed MS-CNN** | 0.92 | 0.867 | 0.89 | 0.95 | 0.94 | 0.91 | 0.94 | 0.92 | 0.95 | 0.92 |

**Figure 13:**
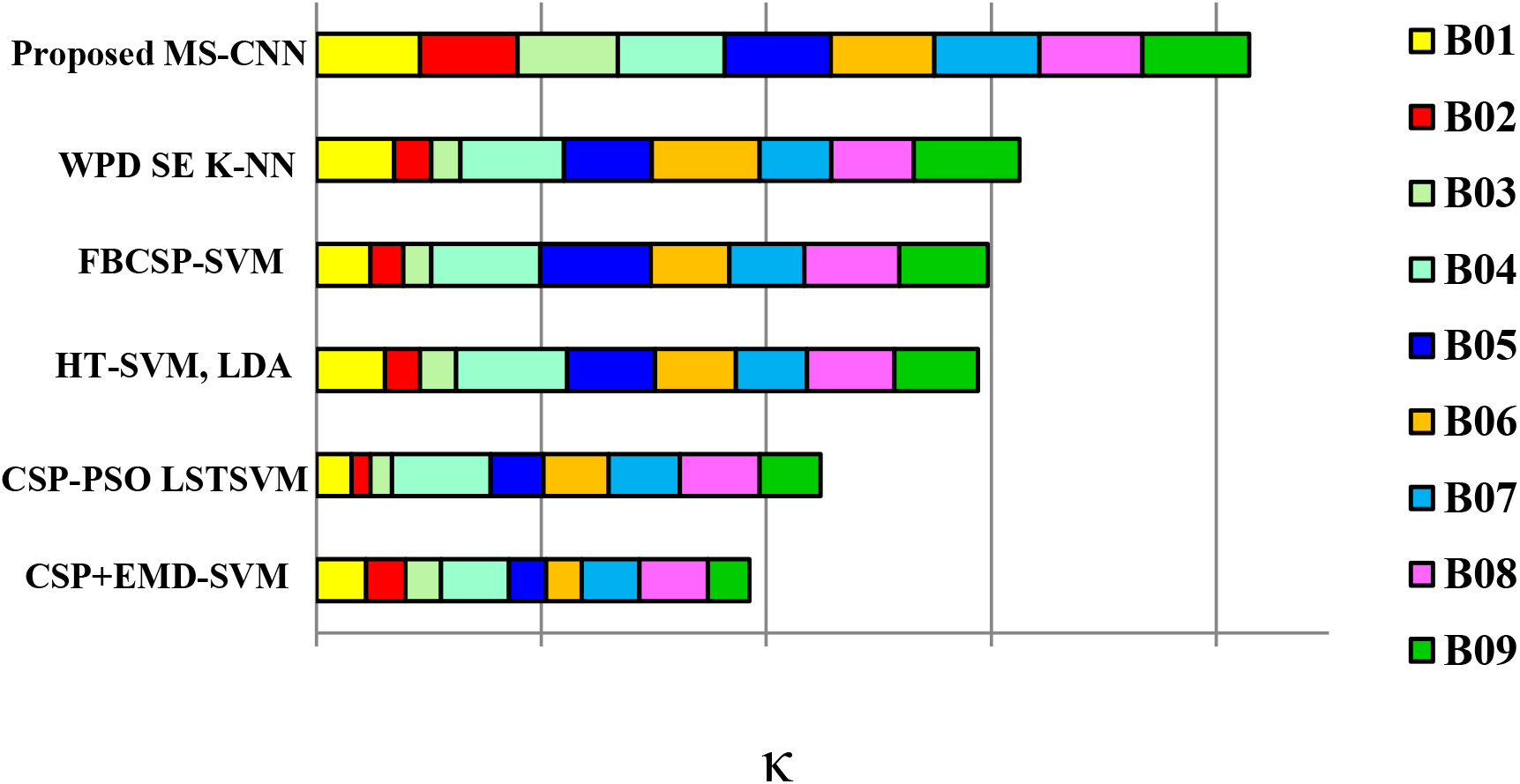
Comparison of *κ* values for each subject between the proposed model (MS-CNN) and other state-of-the-art ML models considering diffrent feature extraction methods and classification algorithms.

### 6.6 Comparison with different state-of-the-art DL models

In this section, the accuracy of several state-of-the-art advanced DL models such as separated channel convolutional neural network (SCCNN)[25], deep transfer CNN (DTCNN) [29], CNN and stacked autoencoders (CNN+SAE) [14], 1-D multi-scale CNN (1DMSCNN) [30], frequential deep belief network (FDBN) [27], and hybrid-scale CNN (HS-CNN) [32] have been compared with the proposed model as detailed in Table 7. It can be seen that the proposed MS-CNN model attains the highest average classification accuracy of 93.74% among other DL models. It is noteworthy to mention that the proposed model improves the average classification accuracy of 11.13%, 9.74%, and 6.14% over the recent and more advanced DL models 1DMSCNN, FDBN and HS-CNN, respectively, as shown in Fig. 14. More specifically, MS-CNN demonstrated significant classification accuracy improvement in subject B01 (up to 12.9% increase), B02 (up to 18.1% increase), B06 (up to 7.2% increase), B09 (up to 9.3% increase) which indicates the efficiency and robustness of the proposed model. From the aforementioned comparison of classification accuracy and *κ* value, it can be inferred that the proposed MS-CNN model demonstrates better accuracy than the state-of-the-art different ML and advanced DL algorithms for all the subjects with the lowest SE. The result indicates the capability of subject-independent representation of EEG data and better generalization of the proposed classifier.

**Table 7:** Comparison of accuracy (in %) of different subjects with existing state-of-the-art DL models where bold indicates the best result from the corresponding model.

| DL Model | Subjects |  |  |  |  |  |  |  |  | Avg. |
| --- | --- | --- | --- | --- | --- | --- | --- | --- | --- | --- |
|  | B01 | B02 | B03 | B04 | B05 | B06 | B07 | B08 | B09 |  |
| SCCNN* [25] | - | - | - | - | - | - | - | - | - | 64.00 |
| DTCNN[29] | 72.6 | 60.3 | 66.9 | 91.2 | 80.6 | 70.6 | 73.2 | 77.7 | 71.2 | 73.77 |
| CNN+SAE[14] | 76.0 | 65.8 | 75.3 | 95.3 | 83.0 | 79.5 | 74.5 | 75.3 | 73.3 | 77.6 |
| 1DMSCNN[30] | 80.56 | 65.44 | 65.97 | 99.32 | 89.19 | 86.1 | 81.25 | 88.82 | 86.81 | 82.61 |
| FDBN[27] | 81.0 | 65.0 | 66.0 | 98.0 | 93.0 | 88.0 | 82.0 | 94.0 | 91.0 | 84.0 |
| HS-CNN[32] | 80.5 | 70.6 | 85.6 | 94.6 | 98.3 | 86.6 | 89.6 | 95.6 | 87.4 | 87.6 |
| **Proposed MS-CNN** | **93.4** | **88.7** | **89.1** | **95.3** | **95.1** | **93.8** | **94.7** | **96.8** | **96.7** | **93.74** |

**Figure 14:**
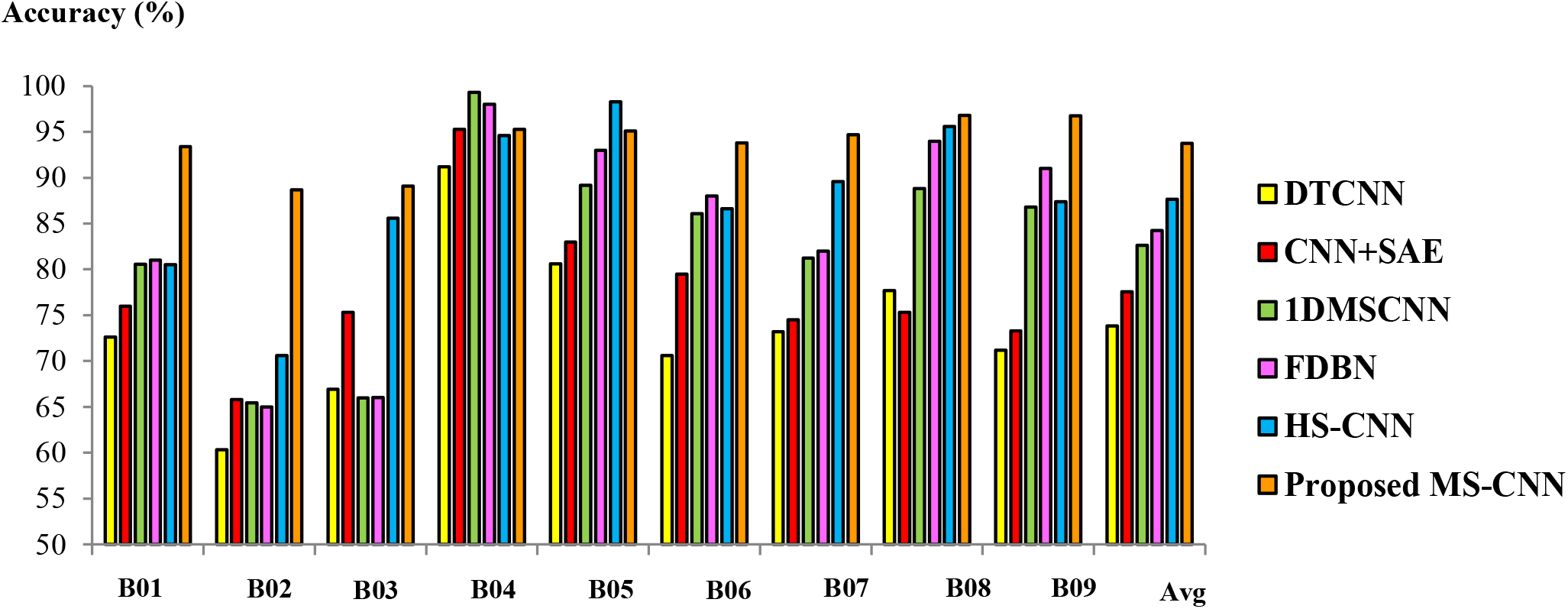
Comparison bar chart of the subject and average accuracy (in %) between proposed model (MS-CNN) and other state-of-the-art DL models for all different subjects.

## 7. Discussion

The current study demonstrates that MS-CNN with DA methods and feature integration techniques significantly improves the accuracy of two class MI-based BCI applications emphasizing the prospect of the current framework in adopting the distinguishable feature of MI-EEG signals. Additionally, the current model reduces the computational burden in MSCB and thus, increase the speed of the classifier. However, the present work focuses on MI-BCI classification tasks for the subject-dependent scenario, where the model has been trained and evaluated on the same subject. The future direction of the current work can be geared toward realizing subject-independent classification [67] where the evaluation phase is independent of the particular subject training class. Additionally, where high-performance person-independent classification is compulsory for the wide application of BCI Systems in the real-world, one possible solution to achieving the goal is to build a personalized model with transfer learning. An efficient transfer model can adopt a transductive parameter to construct an individual classifier which can be further extended to an adaptive-based transfer learning classifier [68]. Another direction could be the implementation of unsupervised or semi-supervised learning [69] to circumvent expensive and time-consuming manual labeling in unsupervised learning to perform classification tasks in abundant class labels for a wide range of MI-BCI scenarios. Moreover, in future work, classification accuracy can be further improved by integrating long short-term memory (LSTM) recurrent neural network (RNN) architecture [70] or self-attention based transformer [71] for extracting semantic temporal-spatial feature of EEG signal and expend the proposed framework for classifying multi-class MI for different BCI applications. Here, some important future research directions, in particular, geared toward the applications of BCI in the healthcare community have been acknowledged. Firstly, the current framework can be utilized in medical and health care, where the deep learning-based BCI systems predominantly work on the detection and diagnosis of mental diseases such as sleeping disorders, Alzheimer’s Disease, epileptic seizure, and other disorders. The MS-CNN model can be widely adopted for its feature engineering and real-time classification for spontaneous EEG stream-based neurodegenerative diseases such as Parkinson’s disease [72]. Moreover, the current model can be suitable for classifying AD based on spontaneous EEG [73] and diagnosis of an epileptic seizure. In such scenarios, a hybrid model containing recurrent neural network (RNN) architecture attached to the MS-CNN model considering tempo-spatial feature extraction can be utilized in seizure diagnosis [74, 75]. Additionally, different mental diseases such as depression [76], Interictal Epileptic Discharge (IED) [77], schizophrenia [78], Creutzfeldt-Jakob disease (CJD) [79], and Mild Cognitive Impairment (MCI) [80] can be detected employing the current BCI deep learning model. Furthermore, the current framework can be extended as a more reliable and robust MI-based real-time brain signal based communications applications such as robotic control [5, 6, 7], P300 speller [81], rehabilitation of neuromotor disorders [4], text entry speech communication [8, 9], cognitive load measurement [82], gaming [2, 3] etc. The current model can be extended to various material modeling [83, 84, 85, 86, 87, 88, 89, 90]

## 8. Conclusion

Summarizing, in this study, a multi-scale convolutional neural network has been designed for EEG-based MI classification. The multi-scale convolution block consisting of different convolutional kernel sizes in the proposed model can extract semantic features in multiple scales for different frequency bands *δ*, *θ*, *α*, and *β* from original EEG data for the classification purpose. Several intrinsic and user-specific features have been extracted from the original EEG data and integrated into the proposed algorithm to improve the accuracy and performance of the model. Furthermore, various data augmentation methods have been utilized to further improve the accuracy and robustness of the proposed classifier by increasing training EEG data. In order to validate the effectiveness of the framework, the proposed model has been applied to the BCI competition IV-2b dataset. Compared with other existing state-of-the-art algorithms, the classification accuracy of the current algorithm has been significantly improved. The results show that the proposed algorithm can attain high classification accuracy with the characteristic of similar performance among the different subjects. With average accuracy of 93.74%, the current framework demonstrates excellent classification performance and generalization. It improves the average classification accuracy of 11.13%, 9.74%, and 6.14% over the recent and more advanced DL models 1DMSCNN, FDBN and HS-CNN (with up to 18.1% increase of accuracy in the subject-specific case), respectively. The proposed model can extract more effective features from EEG signals and can be used to design the efficient and accurate real-time MI-based BCI framework.

## Footnotes

* Ref. [25] do not state accuracy values for each subject.

